# Higher trophic status leads to more diverse and divergent microeukaryote communities over time in urban lakes from the Greater Paris (France)

**DOI:** 10.1101/2025.10.17.683078

**Authors:** Sébastien Duperron, Pierre Foucault, Amaury Le Vern, Midoli Goto, Charlotte Duval, Benjamin Marie, Sahima Hamlaoui, Sébastien Halary, Dominique Lamy, Emilie Lance, Marc Trousselier, Cécile Bernard, Ludwig Jardillier, Julie Leloup

**Affiliations:** Muséum National d’Histoire Naturelle, UMR 7245 CNRS-MNHN, Molécules de Communication et Adaptation des Microorganismes (MCAM), Paris, France; Sorbonne Université, UMR 7618 CNRS-INRA-IRD-Univ. Paris Cité-UPEC, Institut d’Écologie et des Sciences de l’Environnement de Paris (iEES-Paris), Paris, France; Marine Biodiversity, Exploitation and Conservation (MARBEC), Univ. Montpellier-CNRS-Ifremer-IRD, Montpellier, France; Université Paris-Saclay, CNRS, AgroParisTech, Écologie Société et Évolution, 91190 Gif-sur-Yvette, France; Université de Reims, UMR-I 02, Stress environnementaux et biosurveillance des milieux aquatiques (SEBIO), Reims, France

**Author notes:** **Data availability statement** All 18S rRNA gene amplicon sequencing raw reads were deposited into the Sequence Read Archive (SRA) database under the BioProject PRJNA1086840 (see Table S1 for individual sample SRA accession numbers). Scripts available at https://github.com/PierreFoucault/Greater-Paris-microEukaryotes. **Authorship contribution statement** PF: Investigation, Data curation, Formal analysis, Writing – original draft. ALV: Investigation, Data curation, Formal analysis. CD, MG, DL, EL: Investigation. SH, BM, SH, MT, CB: Conceptualization, Investigation, Writing - Review & Editing. JL, SD, LJ: Conceptualization, Investigation, Writing – original draft, Supervision, Project administration, Funding acquisition. **Funding statement** This work, as well as PF and MG grants were funded by the Agence Nationale de la Recherche project COM2LIFE (ANR-20-CE32-0006). ALV was funded by an MNHN. **Competing interest disclosure** The authors declare that they have no known competing financial interests or personal relationships that could have appeared to influence the work reported in this paper.

**Keywords:** eutrophication, freshwater, phytoplankton, protists, time-series, trophic mode, microbial networks

## Abstract

Effects of trophic status on lake microeukaryote community dynamics remain underexplored. Spanning oligotrophic to hypereutrophic conditions, peri-urban lakes located in the Greater Paris region (France) offer unique opportunities to compare these dynamics along a eutrophication gradient. Here, community composition was characterized using 18S rRNA gene metabarcoding, and analyzed in the context of environmental parameters and chlorophyll a-derived trophic status throughout an 18-month sampling period. Microeukaryote assemblages were dominated by Cryptophyceae, Spirotrichea, Chrysophyceae, and Chlorophyceae, with mixotrophic and heterotrophic taxa dominating across all lakes. Taxa richness peaked at intermediate trophic levels, consistent with the intermediate disturbance hypothesis. Beta-diversity and network analyses revealed increasing community modularity, reduced connectivity, and enhanced temporal variability with higher trophic status. In lakes reaching the hypereutrophic status, communities diverged progressively over time, suggesting the onset of regime shifts. Conversely, oligo-to mesotrophic lakes maintained more connected and stable assemblages. These findings demonstrate that eutrophication fosters more diverse and increasingly divergent microeukaryote communities, underscoring its role as a central driver of microbial community restructuring in urban freshwater systems.

## Introduction

Of the 300 million lakes on Earth, approximately 90% are considered small water bodies (less than 1 km^2^; Downing et al. 2006; Pi et al. 2022). Small lakes and ponds harbor and sustain a large biodiversity, play key roles in the continental carbon cycle, at the interface between terrestrial ecosystems and the atmosphere, and act as final collection points for watersheds (Pi et al. 2022; Strayer et Dudgeon 2010). Additionally, they exhibit rapid responses to localized anthropogenic disturbances and are considered as suitable environmental sentinels for assessing the overall health of the surrounding ecosystem (Williamson et al. 2009). Lakes also offer a wide range of ecosystem services including essential supplies of drinking water, crop irrigation, and opportunities for recreational activities. Over the past decades, lakes have experienced significant impacts from human activities, invasive species, increasing surface temperatures and heatwaves, and are considered under threat (O’Reilly et al. 2015; Reid et al. 2019; Williamson et al. 2009; Woolway et al. 2022). These pressures, combined with natural variations, have exacerbated eutrophication, resulting in widespread disruptions in ecosystem functioning, including increased microalgal and cyanobacterial blooms, deoxygenation and salinization (Dugan et al. 2020; Jane et al. 2021). This is particularly pronounced in smaller lakes which are fragmented habitats with limited buffering capacities, and contribute to relatively high amounts of carbon emissions relative to their area (Holgerson et Raymond 2016; Mitsi et al. 2023). Numerous studies have investigated the impact of anthropogenic disturbances on lake prokaryotic communities, identifying global trends, such as highly variable taxa compositions especially under the influence of temperature and nutrient contents (Foucault et al. 2025; Garner et al. 2023; Shen et al. 2019). However, the main ecological functions overall tend to be conserved due to high functional redundancy among taxa (Allison et Martiny 2008; Foucault et al. 2025; Louca et al. 2018).

Lakes also harbor a rich, diverse and abundant community of mainly microbial eukaryotes (Debroas et al. 2017; Escalas et al. 2019; Simon et al. 2015). These have been less investigated despite their major roles in ecosystem functioning, as primary producers, predators, parasites and recyclers of organic matter (Adl et al. 2019; Garner et al. 2022; Itoïz et al. 2022; Monjot et al. 2023; Sommeria-Klein et al. 2021). Recent one-shot explorations at the regional and continental scales have for example revealed how land-use, edaphic conditions and trophic status shape microeukaryotes assemblages (Bock et al. 2020; Garner et al. 2022). On the other hand, even within a limited area, small peri-urban lakes can also display markedly different trophic status owing to the patchy distribution of local anthropic impacts. These offer a unique opportunity to test the impact of trophic status on community compositions by minimizing the effect of confounding factors, and facilitate time-series sampling to evaluate community composition variability. With hundreds of mostly small lakes and reservoirs at the scale of the Île-de-France region (Catherine et al. 2008), Paris is an exceptional natural laboratory to explore these relationships. A previous study pointed out that trophic status was a main driver of bacterial community structure during a summer period, that the functional potential was stable and mostly shared among lakes, and that community heterogeneity increased with higher trophic levels (Foucault et al. 2025).

In the present study, we determined the effect of eutrophication levels on the composition of microeukaryotes communities and their associated trophic mode (phototroph, mixotroph or heterotroph) based on an 18-month survey of communities compositions on nine lakes displaying different levels of eutrophication, all located within 70-km distance around Paris. We hypothesize that i) the eutrophication level has a limited impact on the microeukaryotes and that their temporal variations are comparable because of the close proximity among lakes, ii) that community variability will decrease under higher eutrophication levels owing to the dominance of a limited set of taxa; and that iii) different trophic groups respond differently, with phototrophs being more sensitive than mixotrophs and heterotrophs to local conditions.

## Material and methods

### Sampling

Nine peri-urban lakes were monitored, namely Jablines (JAB), Vaires-sur Marne (VSM), Cergy large (CER-L), Cergy small (CER-S), Créteil (CRE), Bois-le-Roi (BLR), La Grande Paroisse (LGP), Champs-sur-Marne (CSM), and Verneuil-sur-Seine (VSS). Lakes are located within a ∼70 km radius around Paris (France; Fig. S1 and Table S1 for coordinates) and were chosen owing to their similar area and depth (7.3-91.0 ha, 3.5-10 m). All are former sand and gravel quarries converted into leisure centers between the 1960s and the 1980s (Catherine et al. 2008; 2010; Escalas et al. 2019). Lakes were sampled monthly from June 2021 to December 2022. Lack of access prevented sampling on all lakes in December 2021, and at CER-L in June 2021. Overall, a total of 18 time-points were thus sampled from each lake (17 for CER-L).

For each lake and date, the water column was sampled at three mid-lake locations (labeled W1, W2 and W3) to account for spatial heterogeneity. For each water column, 5 L were sampled using a Niskin bottle (WILDCO, USA) at three different depths (∼0.5 m below surface, mid-depth and ∼0.5 m above the lake bottom) and pooled together in equal volumes (depth-integrated sample). Samples were processed on-site within one-hour post-recovery. A total of 483 samples were collected and analyzed.

### Chlorophyll *a* concentration and phytoplankton composition

The chlorophyll *a* (Chl*a*) concentration was measured as a proxy of phytoplankton biomass from 500 mL filtered raw-water (0.7-µm, GF/C, Whatman, UK) in triplicate (Table S2), by spectrophotometry (Cary 60 UV-Vis, Agilent, USA, Yéprémian et al. 2016). The trophic status of the lakes at each date was defined based on Chl*a* concentration ranges from the Carlson’s trophic state index (Carlson 1977): oligotrophic (< 2.6 *µ*g.L^-1^), mesotrophic (2.6 - 7.3 *µ*g.L^-1^), eutrophic (7.3 - 56 *µ*g.L^-1^) and hypereutrophic (> 56 *µ*g.L^-1^; Fig. 1). Phytoplankton taxa were identified based on morphology from lugol-fixed unfiltered water under an inverted microscope (NIKON Eclipse TS100, Japan), using the Utermöhl method (AFNOR 15204 standard, 200 to 400 individuals). The biovolume of each taxa was estimated by multiplying its cell counts by its associated cell biovolume values based on previous reports (Table S3; Escalas et al. 2019; Maloufi et al. 2016). For taxa that were not in these reports, cell biovolumes were extracted from the 2017 HELCOM Phytoplankton Expert Group database (Olenina 2012).

**Fig. 1:**
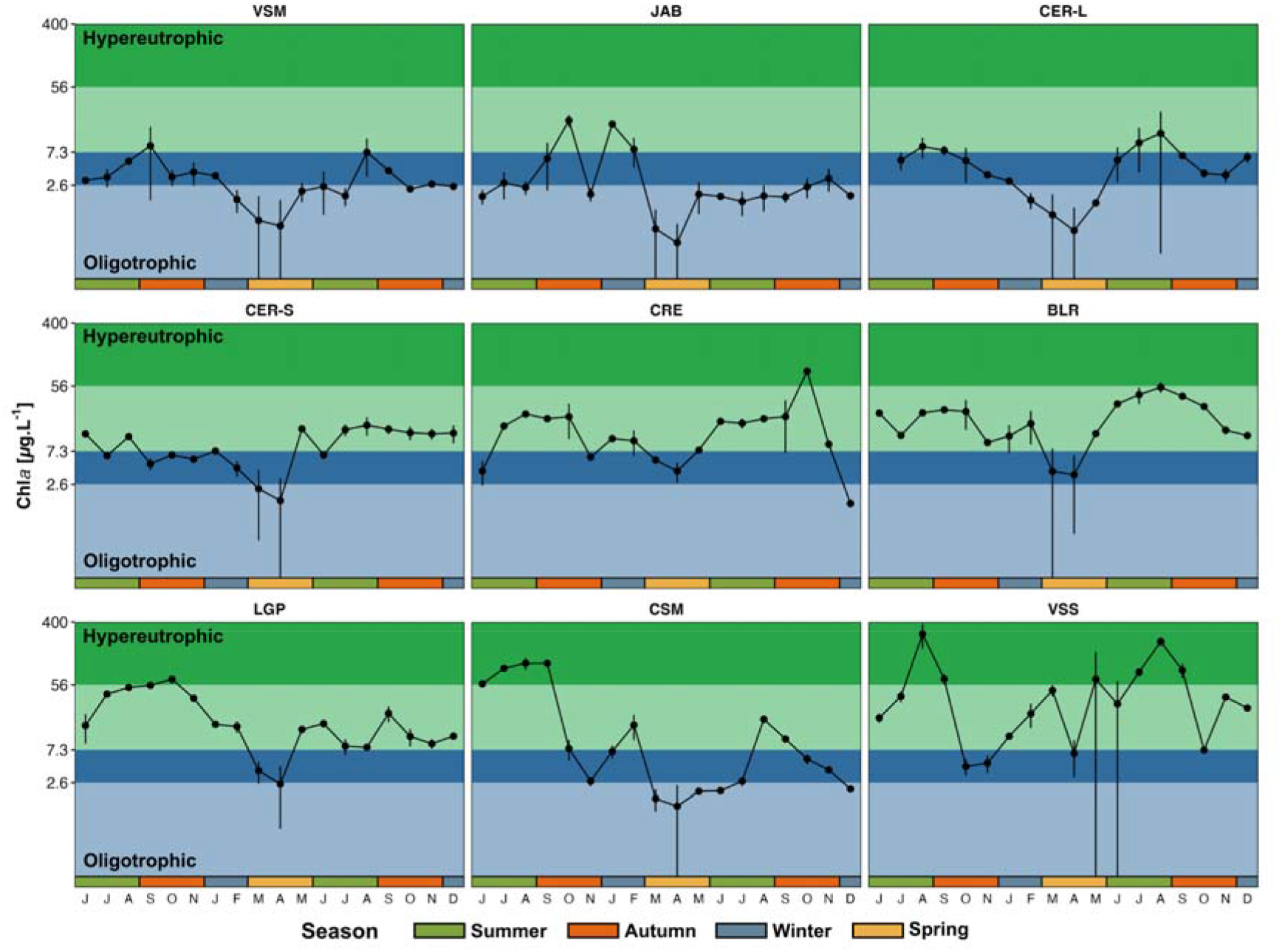
Temporal dynamics of Chl*a* concentrations. Average Chl*a* levels (n = 3 per month, except for VSM in July 2021 n=2) from June 2021 to December 2022. Background colors correspond to Carlson’s TSI trophic states: oligotrophic (light blue), mesotrophic (dark blue), eutrophic (light green), hypereutrophic (dark green, see material and methods). The y-axis is log-scaled, the x-axis corresponds to the month (*i*.*e*., initial of each month, and color bars correspond to seasons).

### Water physico-chemical parameters and trophic status classification

Temperature and pH were measured on shore (KS-2 MultiLine© probe, WTW, USA). Raw water samples were pre-filtered on 50-µm mesh to remove large particles (*e*.*g*., leaves, larvae, largest microbial eukaryotes…). Then, water was filtered onto 0.22-µm PES membranes (Millipore Express, Germany). Eluates were collected in duplicate (12 mL) in polyethylene tubes for nutrient analyses, with an acidification step (three droplets of 3% HNO_3_ solution) for orthophosphate analysis (PO_4_^3-^ ions). Dissolved mineral nitrogen (NH_4_^+^, NO_3_^-^ and NO_2_^-^ ions) and PO_4_^3-^ concentrations were determined as described by Holmes et al. (1999). For total particulate carbon and nitrogen concentration (TPC and TPC), 1 L of raw-water was filtered onto 0.3-µm pre-combusted filters in duplicates (Sterlitec, USA). Filters and eluates were stored at -20°C. Total particulate carbon and nitrogen concentration were determined using a CHN Elemental Analyzer (NA1500 Series 2, Fisons, UK) and normalized by sampled volume (Table S4).

A PCA was performed on water environmental parameters except Chl*a* concentration (T, pH, TPC, TPN, PO_4_^3-^, NH_4_ ^+^, NO_3_ ^-^+NO_2_ ^-^; Fig. S2A; Table S4) revealing that samples were segregated samples by pH and T on the first axis (33.6% expl. var.; cos^2^ 0.43 and 0.42; Fig. S2B) while axis (28.6% expl. var.) was correlated to nutrients (TPC, TPN, PO_4_ ^3-^, NH_4_ ^+^; cos^2^ 0.55, 0.5, 0.48 and 0.43; Fig. S2B). Coordinates on both PCA axis were correlated to Chl*a* concentrations (SPEARMAN, *p*□<□0.001, ρ 0.4 and 0.69; Fig. S2C; Table S5).

### Nucleic acids extraction

For each depth-integrated water column, 150 to 2,000 mL (depending on the lake and the month) of pre-filtered water were filtered onto 0.22-µm PES membranes. Filters were flash-frozen in liquid nitrogen. Total DNA was extracted (483 filtered water samples) using the PowerLyzer PowerSoil DNA extraction kit (QIAGEN, Germany), after a bead-beating step (FastPrep-24 5G, MP Biomedical): 5 × 30s cycles (8 m.s^-1^). Two extraction-blanks and two positive controls were incorporated into the sequencing analyses. Twelve samples were re-sequenced during the second Illumina run to intercalibrate the two sequencing runs.

### 18S rRNA-encoding gene amplicon sequence analysis

The V4 hypervariable region of the eukaryotic 18S rRNA-encoding gene was amplified using primers EUK581-F (5’-GCAGTTAAAAAGCTCGTAGT -3’) and EUK1134-R (5’-TTTAAGTTTCAGCCTTGCG -3’) (Carnegie et al. 2003; Pawlowski et al. 2012). These primers are reportedly biased against metazoans and fungi (Bower et al. 2004). The following program was used: initial denaturation (94 °C, 3 min); 35 cycles (94 °C, 45 s; 55 °C, 60 s; 72 °C, 90 s); final elongation (72 °C, 10 min). Amplicons were sequenced on an Illumina MiSeq 300×2 bp platform (GenoToul, France). A total of 31,200,822 raw reads was obtained.

Sequences were analyzed using the QIIME2 pipeline (version 2024.5; Bolyen et al. 2019). Amplicon Sequence Variants (ASVs) were obtained with the DADA2 algorithm (v1.30.0): reverse reads were trimmed at 245 bp due to lower phred-scores, while the forward read was not trimmed. The expected error rate was set at 3, minimum overlap to 4bp and chimeras were discarded. ASVs were then affiliated taxonomically using the *sklearn* algorithm trained on the PR^2^ database (Guillou et al. 2013) (5.1.0ssu). ASVs affiliated to Bacteria, Metazoa, Mitochondria, Chloroplast, Streptophyta, plastids, nucleomorphs or unassigned were discarded. The remaining dataset contained 14,981,309 high-quality assembled non-chimeric eukaryote reads (31,017 ±13,420 per sample). Rarefaction was performed at 12,287 reads (lowest appropriate sample sequencing depth, after removing four samples, removing samples CER-S_P_W3, CER-L_B_W1, VSM_O_W1 and CSM_N_W1; Table S1). Sample VSM_B_W1 was also discarded from all analyses based on the aberrant measured Chla concentration (Table S2). The analysis yielded 19,714 unique ASVs. Out of these, 19,003 ASVs (96.33%) were considered as ‘rare’, *i*.*e*. their relative abundance never exceeded a 1% abundance threshold in any of the samples, leaving 725 ASVs as abundant (*i*.*e*. their abundance exceeded 1% of the reads in at least one sample). The whole dataset (all ASVs) was used in all analyses except for network analyses. Trophic modes, namely phototroph, mixotroph, phagotroph and parasite, were assigned at the 4^th^ taxonomic rank (PR^2^ Class; Table S6) based on the PR^2^ database (version 5.1.0; Guillou et al. 2013) and additional literature (Adl et al. 2019; Monjot et al. 2023; Singer et al. 2021; Sommeria-Klein et al. 2021).

### Statistical analysis

Analyses were run on Rstudio (v4.4.3, The R Core team 2022). The Chl*a* concentrations were computed by lake and month. The correlation between the PCA axis and the Chl*a* values was assessed by a Spearman correlation test (Rho coefficient (ρ), cor.test, Stats Rbase package v4.4.3).

Alpha-diversity indexes (ASV richness and Shannon diversity) were computed using Phyloseq (v1.50.0; McMurdie et Holmes 2013). Principal Coordinate Analyses (PCoA) were performed based on Bray-Curtis (BC) distances using Vegan (Oksanen et al. 2022). The explanatory power of the ‘lake identity’, ‘season’ and their interaction term were tested using PERMANOVA (*adonis2* from Vegan).

A time lag analysis (TLA) was run on each lake based on date-to-date pairwise Bray-Curtis dissimilarities normalized by the number of days elapsed between respective sampling events. For each dataset, linear and polynomial models (2^nd^ to 5^th^ degree) were fitted. Model fit was evaluated using the Akaike Information Criterion (functions *AIC()* and *lm()* from Lme4 (v1.1-37; (Bates et al. 2015) and LmerTest (v3.1-3; Kuznetsova et al. 2017).

Changes in community composition over time in each lake were estimated using the MOTA (Multivariate Omics Trajectory Analysis) method (Foucault et al. 2022). Briefly, for each lake, “month” centroids are generated by averaging coordinates of the three water columns sampled on a given month. Then, distances between centroids (Euclidian norm) from consecutive time points are summed to compute the length of the total trajectory traveled by monthly centroids during 17 months (starting from July 2021, as CER-L was not sampled in June 2021). The relationship between MOTA trajectory lengths and the 18-month Chl*a* mean of each lake was assessed by a Spearman correlation test (Rho coefficient (ρ), cor.test, Stats Rbase package v4.4.3).

For each lake, a co-occurrence network was built using only the abundant core ASVs of this lake. Abundant core ASVs were defined as that represent at least 1% of the reads in at least one sample from this lake. Networks were built using NetCoMi (v1.2.0; method: SparCC, min. corr. 0.6, Peschel et al. 2021) and igraph (v2.1.4) and network statistics were computed. The percentage of connected nodes assigned to each trophic mode as well as the percentage of modules with one, two, three or the four trophic modes and the trophic mode richness of each module were also calculated.

### Data accessibility

Raw reads were deposited into Sequence Read Archive (SRA, Project PRJNA1086840, see Table S1 for sample accession numbers).

Scripts are available at https://github.com/PierreFoucault/Greater-Paris-microEukaryotes).

## Results

### Temporal variations of trophic status

Temporal variations in trophic status exhibited contrasting patterns across the nine lakes (Fig. 1; Table S2). Lakes VSM, JAB and CER-L maintained a relatively stable low Chl*a* level, within the oligotrophic-to-mesotrophic range (average 3.35 ±2.70, 4.33 ±5.57 and 5.18 ±4.51 *µg*.L^-1^). Lakes CER-S, CRE, BLR and LGP displayed intermediate Chl*a* levels and temporal fluctuations within the mesotrophic-to-eutrophic range. Lakes VSS and CSM showed a larger range of values and reached the hypereutrophic status (0.6 to 134 and 3 to 374 *µg*.L^-1^, respectively). Based on microscopy identification, the phytoplankton community was dominated by eukaryotes throughout the survey period (18-month average: 80.8 ± 28.7%; Fig. S3) in all lakes, except BLR. Four major groups were the most abundant: Diatoms, Cryptophyceae, Dinophyceae and Cyanobacteriota (Fig. S3; Table S7). In the Lake BLR, Cyanobacteriota were dominant (58.8 ± 27.1%; 18-month average).

### Microeukaryote community composition

Microeukaryote communities in the 50-0.22µm size fraction were characterized based on 18S rRNA ASVs metabarcoding, and each ASVs was associated to one of four trophic modes: phototrophs, mixotrophs, phagotrophs and parasites (Table S6). Communities were dominated by our classes: Cryptophyceae (Cryptophyta), Spirotrichea (Ciliophora), Chrysophyceae (Gyrista) and Chlorophyceae (Chlorophyta), whatever the lake, the season or month (Fig. 2 and S4; Table S8). Phagotrophs and parasites represented 30.1 % and 21.4 % of all ASVs; and 30.9 % and 6.2 % of all reads, respectively. Phototrophs accounted for 29.7 % of all ASVs and 23.0 % of all reads. Finally, mixotrophs only accounted for 6.6 % of all ASVs, but represented 39.0 % of all reads. The most prevalent ASV, identified as the mixotroph *Cryptomonas curvata* (Cryptophyta), alone accounted for 18.9 % of total reads in the whole dataset, emphasizing the abundance and ubiquity of certain taxa.

**Fig. 2:**
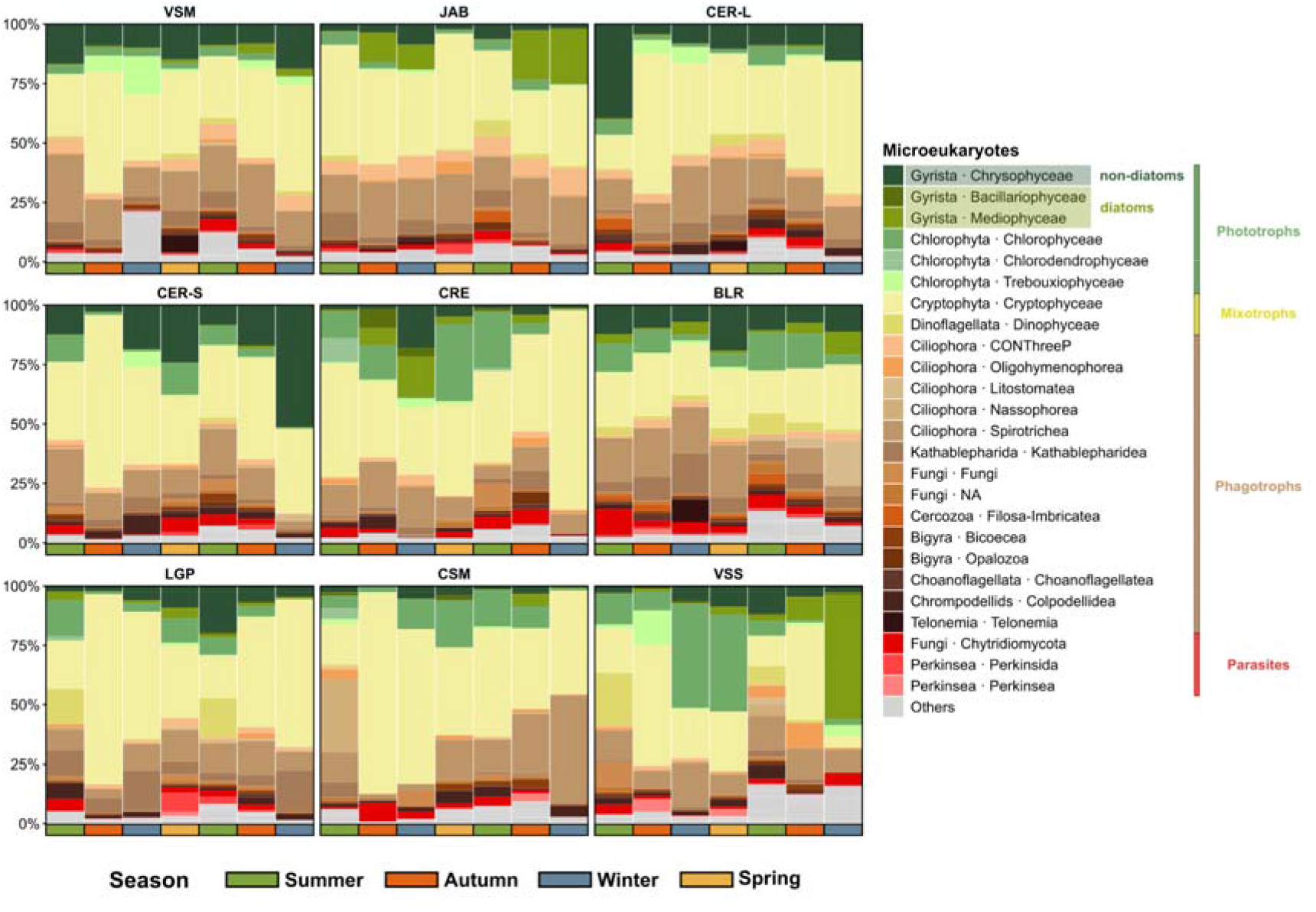
Taxonomic composition of microeukaryote communities. Median proportion of the total ASV reads per season. The 25 most abundant classes are colored according to their potential trophic mode (phototrophs, mixotrophs, phagotrophs and parasites). On the x-axis, the color bars correspond to seasons. Lake panels are ordered according to increasing 18-month averaged Chl*a* concentration (from left to right, then from top to bottom).

All the microeukaryote communities exhibited similar temporal patterns in alpha diversity indexes, with higher ASV richness observed during the summer (Fig. S5). Values ranged from 116 ± 72 ASVs in the winter of 2021 to 304 ± 99 in the summer of 2022 (Fig. S5A). Similar seasonal patterns were observed in the different trophic modes (Fig. S5C-F). No evident trend with the trophic status of the lakes was observed for any of the diversity indexes. Indeed, lakes with the lowest (JAB, oligotrophic-to-mesotrophic) and highest (VSS, eutrophic-to-hypereutrophic) average Chl*a* concentrations both exhibited lower ASV richness than lakes displaying intermediate Chl*a* levels.

Community structure comparisons (Bray-Curtis dissimilarity) indicated, significant segregation by lake (*p* < 0.001 and R^2^ = 0.21), then, to a lower extent, by season (*p* < 0.001 and R^2^ = 0.07; Table S9). However, there was some overlap among most lake’s polygons (Fig. S6). The microeukaryote communities of the different lakes displayed varying temporal dynamics (PERMANOVA; lake:season interaction; *p* < 0.001 and R^2^ = 0.16): a significant effect of trophic status was observed among the communities of the oligotrophic-to-mesotrophic (VSM, JAB and CER-L), meso-eutrophic (CER-S, CRE, BLR, LGP) and the hypereutrophic (CSM and VSS) range for lakes (*p* < 0.001 and R^2^ = 0.08; Table S9). Specifically, the trophic status of lake CSM transitioned over time from hypereutrophic (summer and early autumn of 2021) to mesotrophic status (2022), with occasional shifts to eutrophic and oligotrophic levels (Fig. 1). This major change is reflected on its associated eukaryotic communities compositions, with limited overlap between polygons of summers 2021 and 2022, and no overlap between autumns 2021 and 2022 (Fig. S6). Concerning the lake BLR, its polygons displayed very limited overlap with those from other lakes, indicating different communities (Fig. S6). Besides, the BLR polygons corresponding to each season were very small and largely overlapping with one another compared to all other lakes, suggesting a stable community composition in this lake.

When focusing on mixotroph, phototroph and phagotroph trophic modes, community compositions were also affected by both lake and season (*p* < 0.001, R^2^ values ranging from 0.16 to 0.25 for lakes, 0.04 to 0.08 for season; Fig. S7; Table S9). For parasites, the effect of lake identity was of lesser importance (*p* < 0.001, R^2^ values 0.09 for lakes and 0.03 for season; Fig. S7; Table S9). For all trophic modes the seasonal effect on the community composition varied greatly between lakes (*p* < 0.001, R^2^ values ranging from 0.15 to 0.17 for lake and season interactions; Fig. S7; Table S9).

Overall similar temporal patterns of microeukaryote community dissimilarities were found in all nine lakes based on TLA. The best fit was obtained using polynomial rather than linear models (Fig. 3; Table S10). Month-to-month dissimilarities were highest when comparing communities approximately 6 months apart (opposite seasons) and lowest for those 1 month apart. When looking at intervals up to 1 year, higher dissimilarities were measured in CSM and VSS lakes (high Chl*a* levels), indicating higher variability in community compositions. Additionally, these two lakes were the only ones for which a linear correlation with a significantly positive slope was found (Fig. 3; Table S10). This suggests that microeukaryote communities at CSM and VSS lakes did not return to their initial state after one year, and diverged with time, beyond the seasonal oscillations. For the other lakes, non-significant (*p* > 0.05) or slightly negative slope (JAB lake) were observed.

**Fig. 3:**
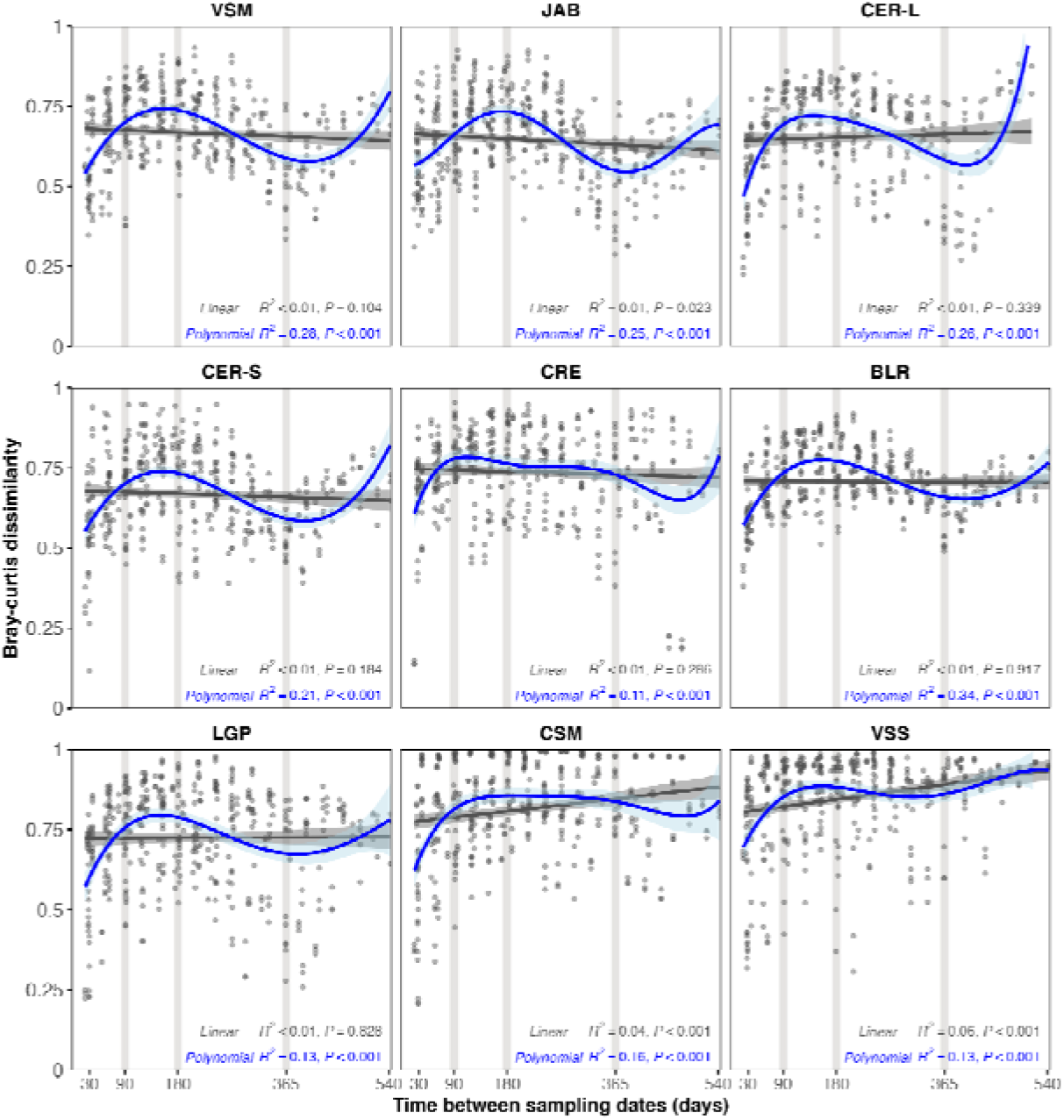
Time-lag analysis of microeukaryote community dissimilarities. Date-to-date Bray-Curtis dissimilarities computed over all time intervals for each lake (from 30 to 540 days). Linear (black) and polynomial (blue) regression are displayed. Significance and rho statistics of each regression are displayed. The 3-month, 6-month, 12-month time lags are highlighted with vertical grey backgrounds. Lake panels are ordered according to increasing 18-month averaged Chl*a* concentration (from left to right, then from top to bottom).

The amount of change in community composition over time was estimated by computing the length of the total trajectory traveled by monthly centroids using the MOTA method. Shortest total lengths were obtained for the oligo-to-mesotrophic VSM, JAB and CER-L lakes, and longer trajectory lengths were observed for other lakes that reached the eutrophic and hypereutrophic trophic status (Fig. 4A). Lake VSS displayed the longest trajectory, again indicating highest variability in community composition. Total trajectory length was correlated to lakes’ 18-month Chl*a* mean concentration (Fig. 4B; Table S5) as well as concentration range (Table S5).

**Fig. 4:**
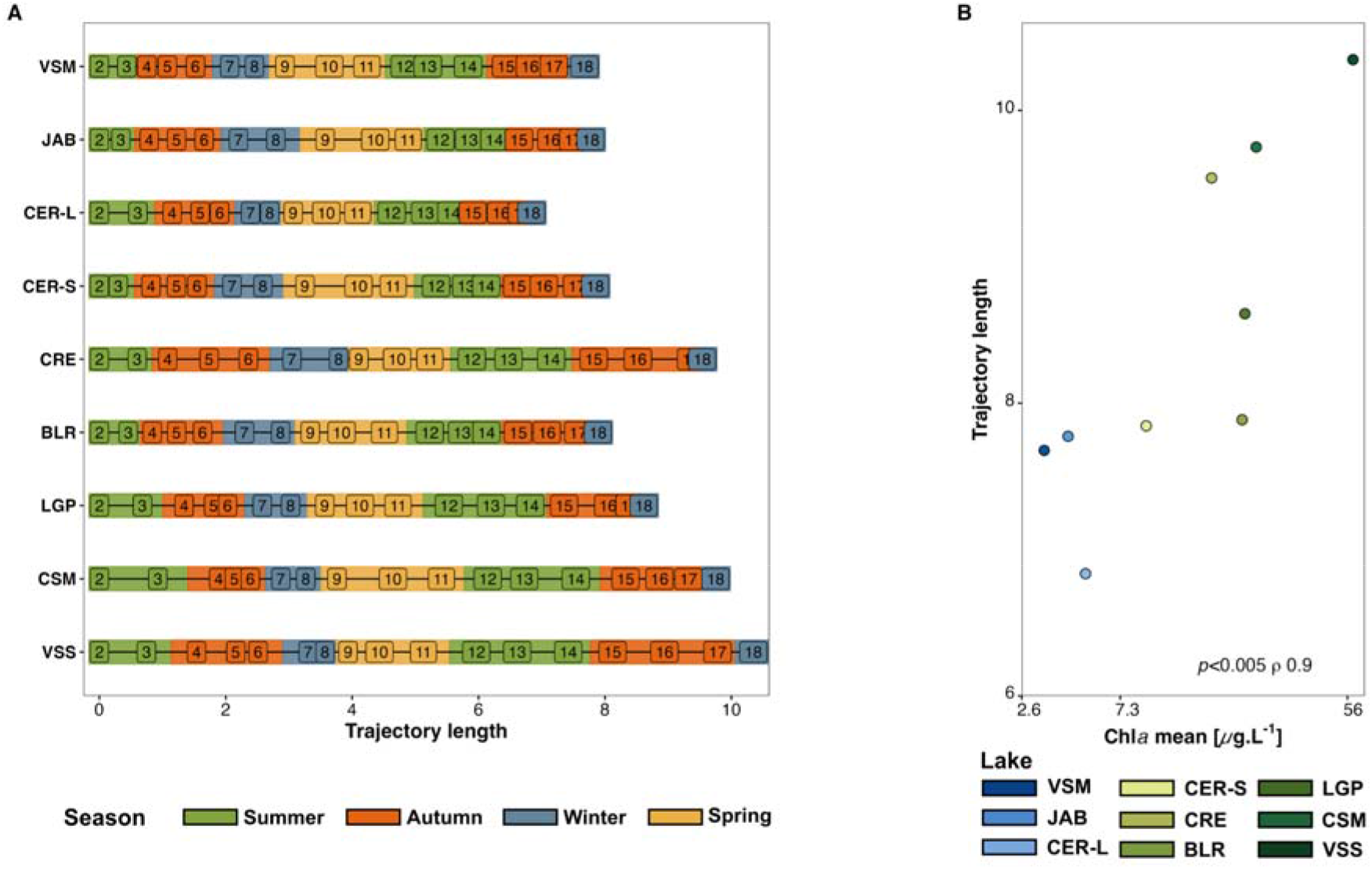
Relationship between MOTA trajectory length and Chl*a* concentrations. **A:** MOTA month-to-month cumulative trajectories lengths of the communities (BC dissimilarity). Trajectories were calculated on the first 56 axes (90.26% of the total explained variance). The number corresponds to the sampling month (*i*.*e*., 2 for January and 12 for December). The lakes (y-axis) are ordered according to increasing 18-month averaged Chl*a* concentration (from top to bottom). **B**: Relationship between the trajectory length in a given lake and its 18-month mean Chl*a* concentration. The x-axis (Chl*a* concentration) was log-scaled. Spearman rho statistics is displayed.

### Co-occurring taxa and network properties

The number of lake-specific core ASVs ranged from 126 (JAB and CRE) to 188 ASVs (VSS; Fig. 5; Table S11). The nine networks contained a similar number of connected nodes (52.67 ±9.71) and modules (*i*.*e*. groups of at least two connected nodes; 12.56 ±2.7; Fig. 5; Table S11). Number of core ASVs, components, clustering coefficient, modularity as well as percentage of positive edges increased with increasing trophic status category from low (VSM, JAB, CER-L) to intermediate (CER-S, CRE, BLR and LGP) and high Chl*a* levels (CSM and VSS; Fig. 5; Table S11). Trophic richness per module increased slightly (Table S11). Overall these results indicate that lakes of higher trophic status have more structured networks, which are likely sensitive to the removal of certain taxa. On the other hand, the networks of the oligotrophic-to-mesotrophic lakes (VSM, JAB and CER-L) displayed a higher number of edges compared to others (Fig. 5; Table S11). Moreover, their largest connected component contained a higher fraction of the network nodes: 26.8 versus 20.1 and 6.2% for the other two categories (Fig. 5; Table S11), indicative of more connected members of the community. Regarding trophic modes, all lakes had comparable percentages of connected nodes affiliated to a given trophic mode: phototrophs (avg. 38.0 ±7.5%), phagotrophs (33.63 ±7.6%), mixotrophs (18.7 ±3.9%), then parasites (9.6. ±3.1%; Table S11). The network of lake BLR showed notable differences compared to others; it displayed the most connected nodes, largest connected component (LLC, 42.4%), highest percentage of negative edges (39.2%), highest module mean trophic richness (2.56) and percentage of modules with the four trophic modes (22.2%), and lowest clustering coefficient (0.41; Fig. 5; Table S11), overall indicating more complex trophic interactions.

**Fig. 5:**
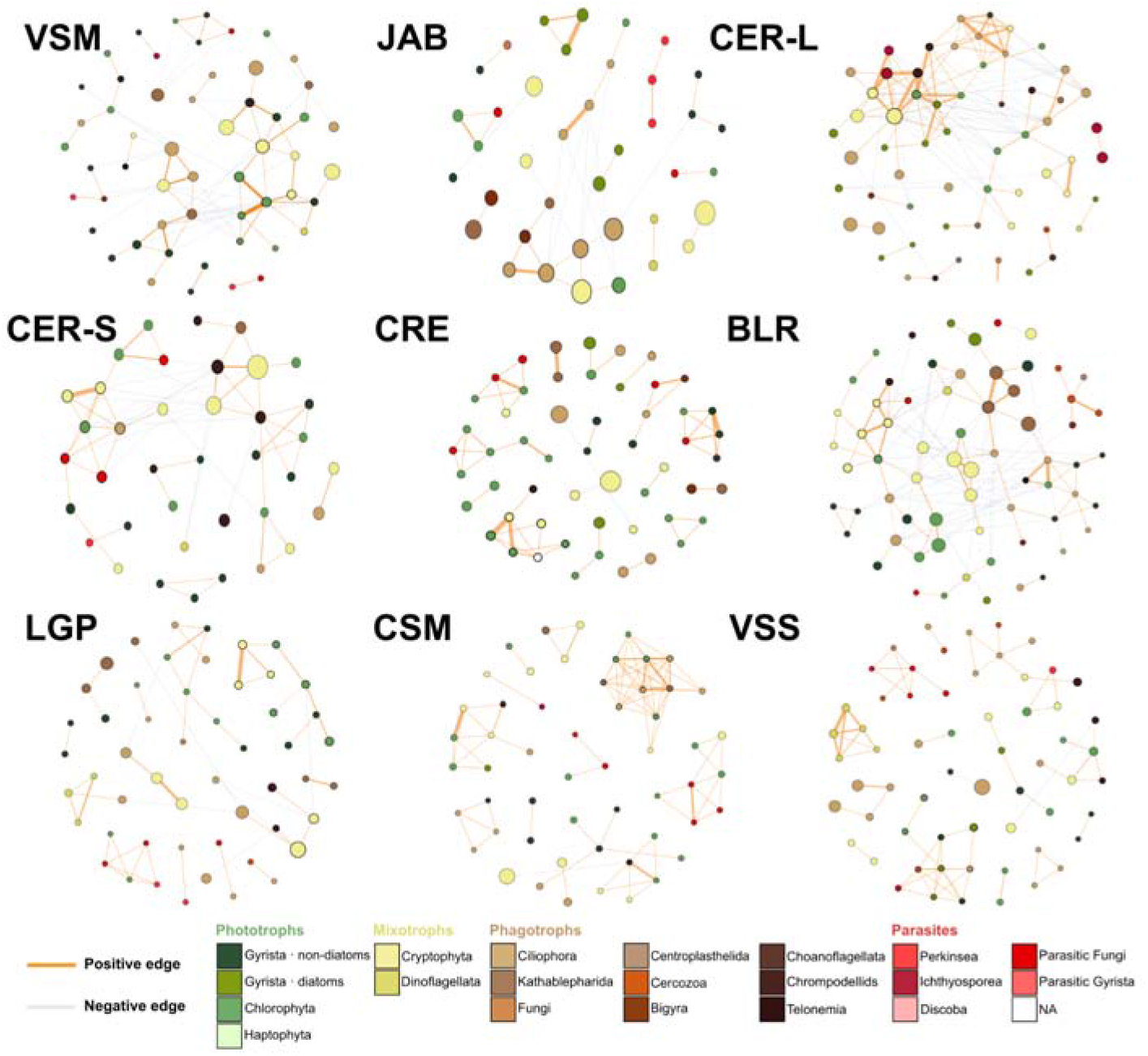
Co-occurrence networks of the microeukaryote communities. Co-occurrence networks based on the lake-specific core ASVs (>1% of the reads in at least one sample). The node diameter indicates mean relative abundance of ASVs and the edge thickness corresponds to the weight of the correlation. The nodes are colored according to the ASV potential trophic mode (phototrophs, mixotrophs, phagotrophs and parasites, see Fig. 2). Lake panels are ordered according to increasing 18-month averaged Chl*a* concentration (from left to right, then from top to bottom). Only connected nodes are shown.

## Discussion

The microeukaryote communities of the nine lakes, as estimated by ASVs, were dominated by Cryptophyceae (Cryptophyta), Spirotrichea (Ciliophora), Chrysophyceae (Gyrista) and Chlorophyceae (Chlorophyta). These results support the abundance and importance of mixotrophs and heterotrophs in lakes, and suggest the presence of a common group of generalists, likely phagotrophs (mixotrophs and heterotrophs) year-round across the geographical area, whatever the trophic status of lakes. In our case, *Cryptomonas curvata* is one such example, being abundant in all lakes and seasons. This is congruent with various studies showing sets of Ciliates, Cryptophyta (including *Cryptomonas*) and Chrysophyceae (including *Dinobryon*) ASVs consistently found year-round as core members in both temperate and arctic lakes (Potvin, Rautio, et Lovejoy 2021; Šimek et al. 2023; Simon et al. 2015). In contrast, no shared set of autotrophs ASVs was identified over the nine lakes, indicating greater variability within this trophic group, suggesting higher sensitivity to local conditions. However, this finding could be due to the underrepresentation of some taxa, as many phototrophic taxa are larger than the size-fraction fraction used for amplicon analysis, including members of the Chlorophyta, Gyrista (specifically diatoms, which were particularly more abundant in the phytoplankton counts than in the 18S rRNA amplicon survey) or Dinoflagellata. In the same vein, parasitic taxa were present in low abundance in all lakes, as recently found by Garner et al. (2022) in their large survey of Canadian lakes. Parasites may also be underrepresented, due to their association with organisms larger than the size fraction, such as metazoan and larger phytoplanktonic taxa. Additionally, as the primers used are biased against fungi, some taxa like ecologically-important chytrids that reportedly infect Cyanobacteriota (Bower et al. 2004; Gleason et al. 2008; McKindles et al. 2021) and are abundant in temperate lakes such as Lake Sanabria, Spain (Mitsi et al. 2023), may have been overlooked. Despite these possible limitations, lake-specific patterns were observed in correlation to the trophic status of these lakes.

### Taxa diversity is higher in lakes of intermediate trophic status

Lower species richness was detected for the communities in lakes with both the highest and the lowest Chl*a* levels compared to lakes with intermediate levels. This is consistent with the Intermediate Disturbance Hypothesis, which predicts diversity peaks at intermediate levels of disturbance or environmental stress, while both low and high extremes tend to favor a reduced set of taxa (Weithoff et al. 2001). Indeed, oligotrophic conditions represent an extreme due to nutrient scarcity, whereas hypereutrophic conditions represent another extreme due to the competitive exclusion exerted by a few dominant taxa. Both conditions select for a more limited number of taxa compared to intermediate conditions (Liu et al. 2019). Recently, lakes of higher trophic status were for example reported to display lower diversity of protists in a large-scale survey (Canada, Garner et al. 2022), as well as in a survey over a more limited region (IDF-France, Maloufi et al. 2016; Escalas et al. 2019). In these studies, dominant taxa included bloom-forming phototrophs as well as various heterotrophs benefiting from high nutrient levels and primary production (Huisman et al. 2018; Garner et al. 2022). On the other hand, intermediate trophic status in a lake may favor a more diverse community by providing more diverse niches for organisms.

### Community structuration and divergence are affected by high trophic status

A significant but limited difference in community composition was observed based on the trophic status, especially when comparing the extremes. The maximum difference was found between the oligotrophic-to-mesotrophic lakes JAB, VSM and CER-L and the eutrophic-to-hypereutrophic lakes VSS and CSM. Stronger community structuring in modules was observed in the networks of lakes with higher trophic status, with each module consisting of different taxa, and limited connectivity among them. Such community structuration suggests that these modules may correspond to taxonomically distinct assemblages potentially associated with different ecological functions, which could imply lower functional redundancy and increased niche compartmentalization (Xue et al. 2018). Such a pattern would make higher-trophic-status lakes communities more vulnerable to functional loss if specific modules are affected by environmental disturbances. These features could make the communities of eutrophic-to-hypereutrophic lakes more sensitive to environmental disturbances, consistent with their higher temporal variability. In contrast, networks from oligo-to mesotrophic lakes show more connections among taxa and lower modularity, suggesting less niche segregation, reduced sensitivity to taxa loss, and greater functional redundancy. This pattern is also consistent with the greater temporal stability observed for these communities. However, it should be noted that such stability occurs under comparatively less variable environmental conditions in these lakes. Thus, it does not necessarily imply that these communities would be better able to cope with eventual disturbances than those of lakes VSS and CSM.

Lake BLR appears as an outlier compared to others, owing to its stability in terms of both trophic status and microeukaryote community composition, as well as the dominance of Cyanobacteriota in the phytoplankton. Its associated network is the most diffuse, with the highest number of connections, lowest clustering coefficient, and highest number of trophic modes per module indicating stable communities with a complex network of interactions. In a previous study, prokaryotic communities were also found to be highly stable in lake BLR and different from other lakes (Foucault et al. 2025) suggesting that peculiar local conditions shape both eukaryotic and prokaryotic communities. However, none of the parameters examined in the present study explained the difference with other lakes, and local peculiarities (pedology, type of vegetation, land use, presence of unmeasured contaminants) should be further explored to explain the difference between this and the eight other lakes.

### Stable temporal patterns are detected but hypereutrophic status induce community shifts

Similar temporal patterns are observed in all lakes, with a summer alpha diversity (richness) being around twice that observed in winter, which is not surprising given their close geographical proximity. This trend holds true for all microeukaryotes trophic modes (phototrophs, mixotrophs, phagotrophs and parasites). Higher eukaryote diversity during summer and autumn has often been observed in temperate lakes, as evidenced in the 20 years’ time-series conducted in Lake Mendota (Krinos et al. 2024). Within a one-year interval, community compositions are most different in opposite seasons (winter versus summer, for example), as expected under temperate climates (Simon et al. 2015; David et al. 2021). They tend to be more similar after one year, indicating cyclic variations in community structure, as regularly documented in longer time series (David et al. 2021; Krinos et al. 2024). However, despite overall similar trends, lakes that reach the hypereutrophic status (CSM and VSS) displayed unique features. First, higher variability of communities was observed throughout the 18-month period, not limited to the maximal primary productivity period (*i*.*e*. summer), with higher month-to-month dissimilarities compared to other lakes. This is congruent with the hypothesis formulated by Garner et al. (2022), based on their Canadian survey, that protist communities in productive lakes are less stable over time. Second, community compositions in CSM and VSS lakes also tend not to return to their initial state after one year, instead showing signs of year-to-year divergence, in a way similar to that observed by David et al. in microeukaryotic communities over a 2-year time series on a set of small ponds (2021). This trend is not observed in lakes displaying lower trophic status, suggesting that hyper-eutrophication may ultimately lead to increased community composition drift with time. The hypothesis of a regime shift in the same hypereutrophic CSM and VSS lakes was recently formulated for prokaryotic communities investigated during the summer of 2021 (Foucault et al. 2025). Thus, our results suggest that microeukaryote communities in lakes reaching the hypereutrophic status also start diverging with time. Whether hypereutrophic conditions represent a tipping point beyond which the divergence of microeukaryote communities increases over time will require additional testing for year-to-year drift over a period of several years.

## Conclusion

This study shows that the eutrophication level has a limited impact on microeukaryotes community compositions within the size fraction examined herein, except when the hypereutrophic status is reached. Diversity is maximal at intermediate trophic status, congruent with the Intermediate Disturbance Hypothesis. Mixotrophic and heterotrophic taxa dominate with a few taxa being abundant in all lakes year-round, while phototrophs are less conserved. Lakes reaching the hypereutrophic status show markedly increased variations throughout all seasons, and potential year-to-year drift in community compositions compared to other lakes, suggesting the existence of a tipping point affecting lake community dynamics. Long-term monitoring of lakes spread over a limited area, such as the Ile de France region, provides an opportunity to test the hypothesis of a regime shift and identify the exact conditions that trigger it.

## Acknowledgements

Authors thank the members of the COM2LIFE consortium for their participation in the fieldwork (authors of this work, Siham Mesli, Maëlle Jarno) and especially Haeitz Aloui for the administrative work. We thank the directors of the recreational centers for their monthly access approval. We thank the GENOTOUL sequencing facility.

