## Supplementary Figures for "Higher trophic status leads to more diverse and divergent microeukaryote communities over time in urban lakes from the Greater Paris (France)"

**Supplemental data**

**Table S1: Samples identification and preprocess summary**

**Table S2: Chla concentration dataset (in µg.L^-1^)**

**Table S3.1: Phytoplankton taxonomy, cell count and relative abundance**

**Table S3.2: Phytoplankton taxonomy and associated biovolumes (in µm^3^)**

**Table S4: Physico-chemical parameters dataset**

**Table S5: Summary of Spearman's rank test results**

**Table S6: Microeukaryote taxonomy (PR^2^) and associated trophic mode**

**Table S7: Microeukaryotes taxonomy (PR^2^) and relative abundance by lake and season**

**Table S8: Phytoplankton taxonomy and relative abundance by lake and season**

**Table S9: Summary of permanova results**

**Table S10: Summary of linear and polynomial model regressions results**

**Table S11: Summary of network properties**

**Fig. S1: Location of the lakes within the Île-de-France region (France)**

**Fig. S2: Correlation between Chl*a* concentration values and other water parameters**

**Fig. S3: Temporal dynamic of the phytoplankton community composition**

**Fig. S4: Temporal dynamic of the microeukaryote community composition**

**Fig. S5: Temporal dynamic of the diversity indexes of microeukaryote communities**

**Fig. S6: PCoA plots of the microeukaryote communities**

**Fig. S7: PCoA plots of the microeukaryote communities based on trophic modes**

**Fig. S1: Location of the lakes within the Île-de-France region (France)**

Lakes identifiers are: Jablines (JAB), Vaires-sur Marne (VSM), Cergy large (CER-L), Cergy small (CER-S), Créteil (CRE), Bois-le-Roi (BLR), La Grande Paroisse (LGP), Champs-sur-Marne (CSM), Verneuil-sur-Seine (VSS). GPS coordinates are in Table S1.
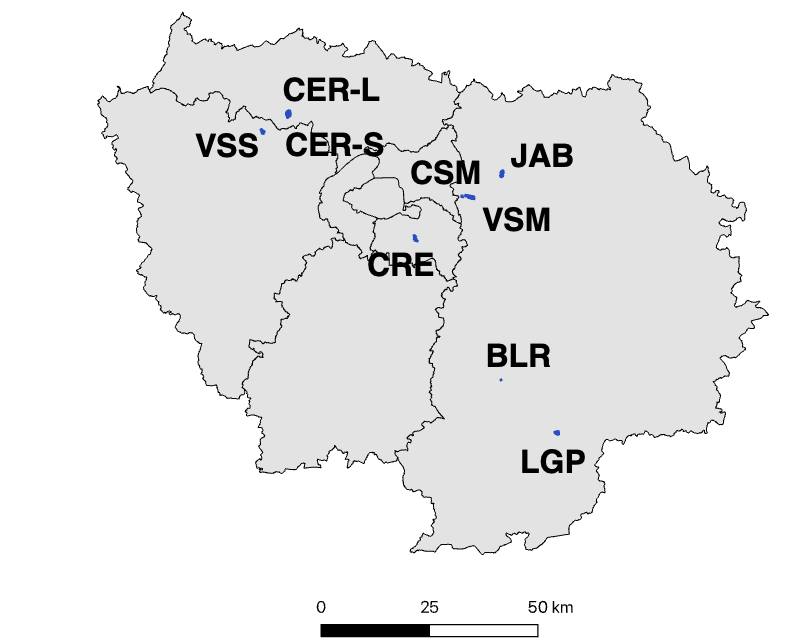

**Fig. S2: Correlation between Chl*a* concentration values and other water parameters.**

**A:** PCA based on water environmental parameters (T, pH, TPC, TPN, PO_4_^3-^, NH_4_^+^, NO_3_^-^+NO_2_^-^) from each lake are displayed in individual panels, with identical axis coordinates, and ordered according to their 18-month averaged Chl*a* concentration (from left to right, then from top to bottom). Seasons are colored and delimited by polygons representing the maximal area delimited by the sample’s coordinates. **B:** Correlation (Cos^2^ values) of each feature with the two first axes of the PCA based on scaled and centered water parameters except Chl*a* concentration. **C**: Relationship between the PC2 axis coordinates and the Chl*a* concentration. The x-axis (Chl*a* concentration) was log-scaled. Spearman rho statistics is displayed.

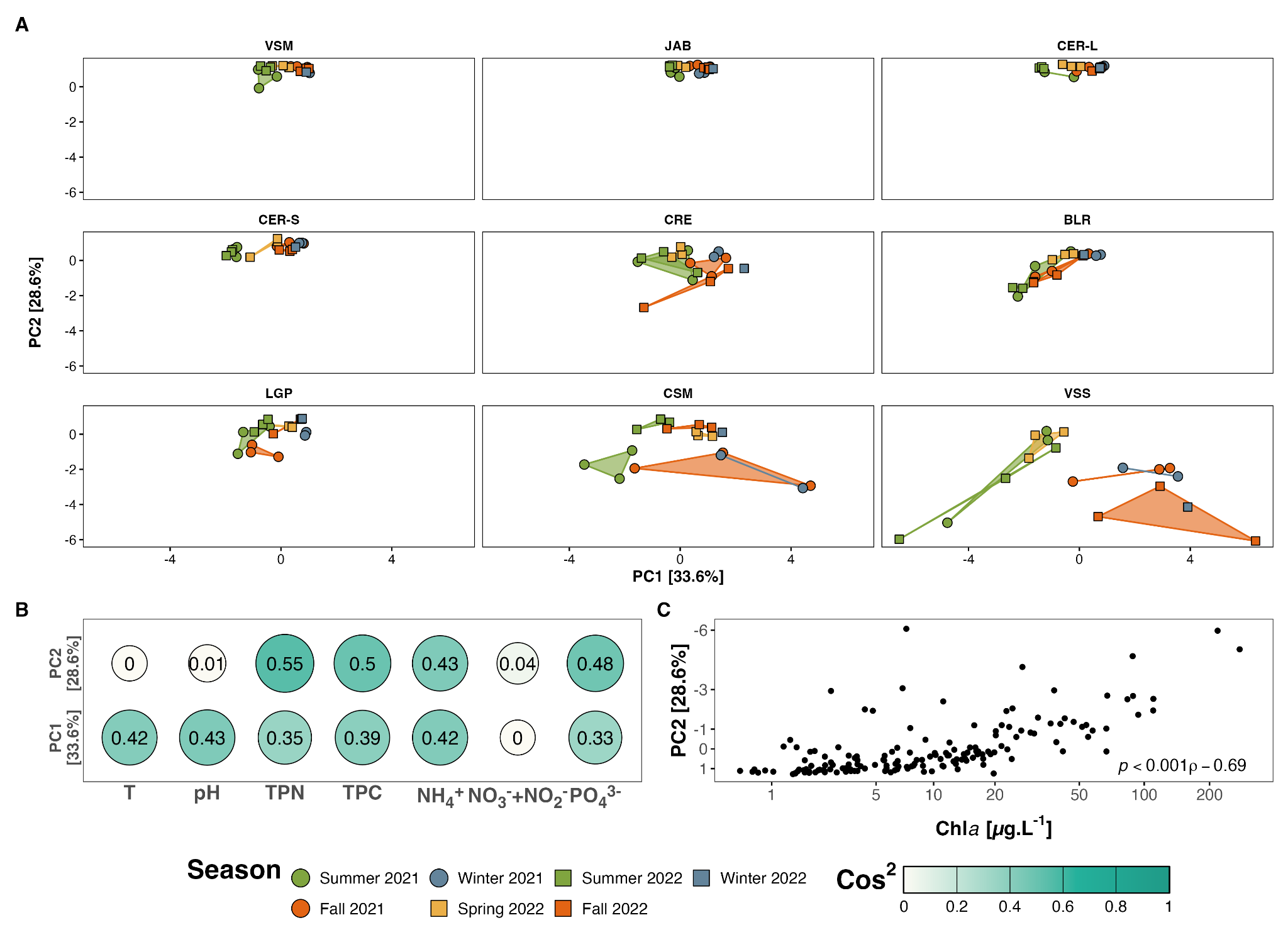

**Fig. S3: Temporal dynamic of the phytoplankton community composition**

Relative biovolume (biovolume-adjusted cell count) of the eukaryote and prokaryote phytoplankton (PR^2^-based Subdivision and Class rank) for each lake as percentage of the total phytoplankton biovolume. Lake panels are ordered according to increasing 18-month averaged Chl*a* concentration (from left to right, then from top to bottom). The x-axis corresponds to the month (*i.e.,* initial of each month) and the color bars correspond to seasons.

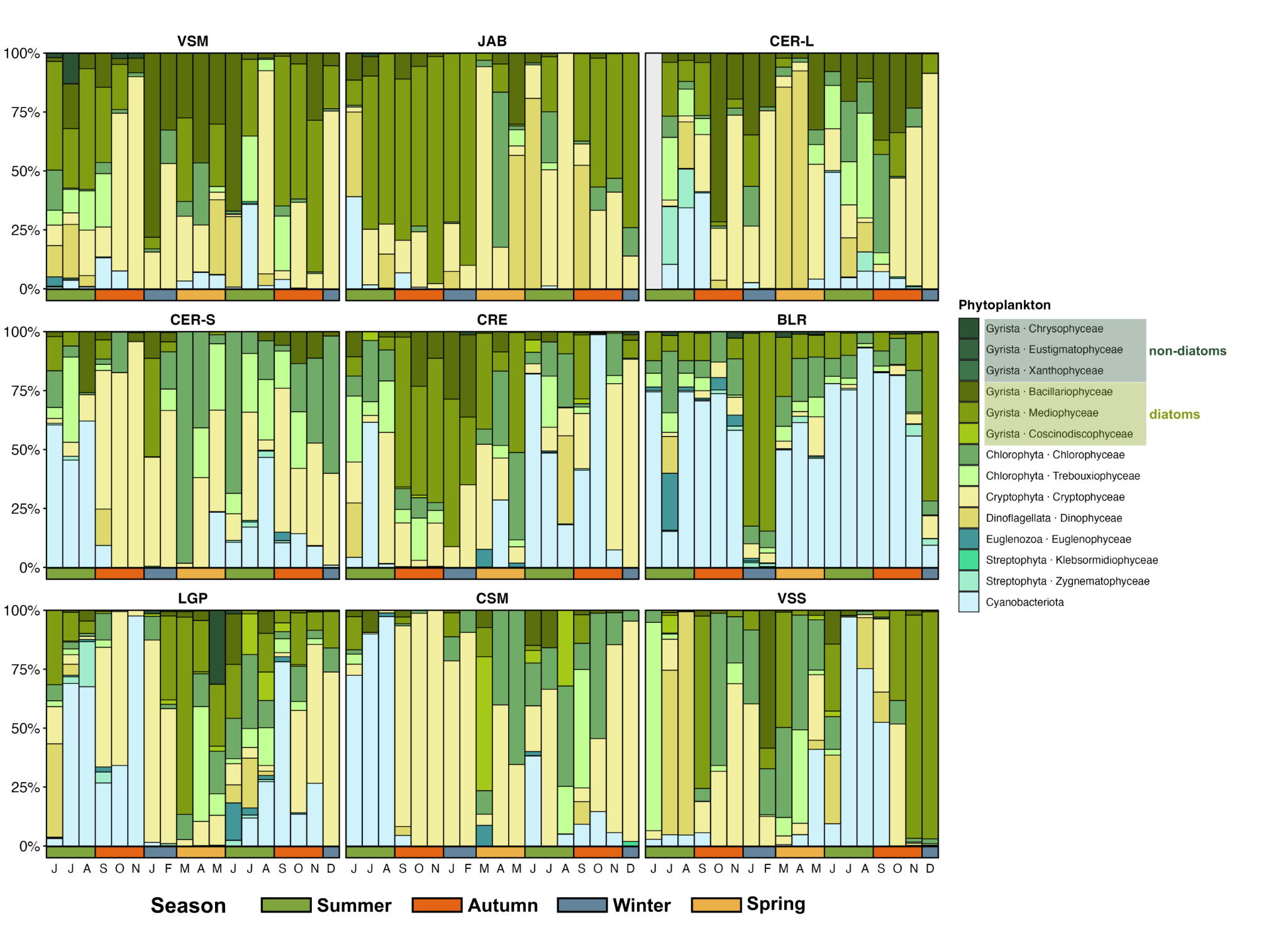

**Fig. S4: Taxonomic composition of microeukaryote community** **composition.**

Median proportion of total ASV reads per month. The 25 most abundant classes are colored according to their potential trophic mode (phototrophs, mixotrophs, phagotrophs and parasites). On the x-axis, the letters correspond to the months (June 2021 to December 2022) and the color bars to the seasons. Lake panels are ordered according to increasing 18-month averaged Chl*a* concentration (from left to right, then from top to bottom).

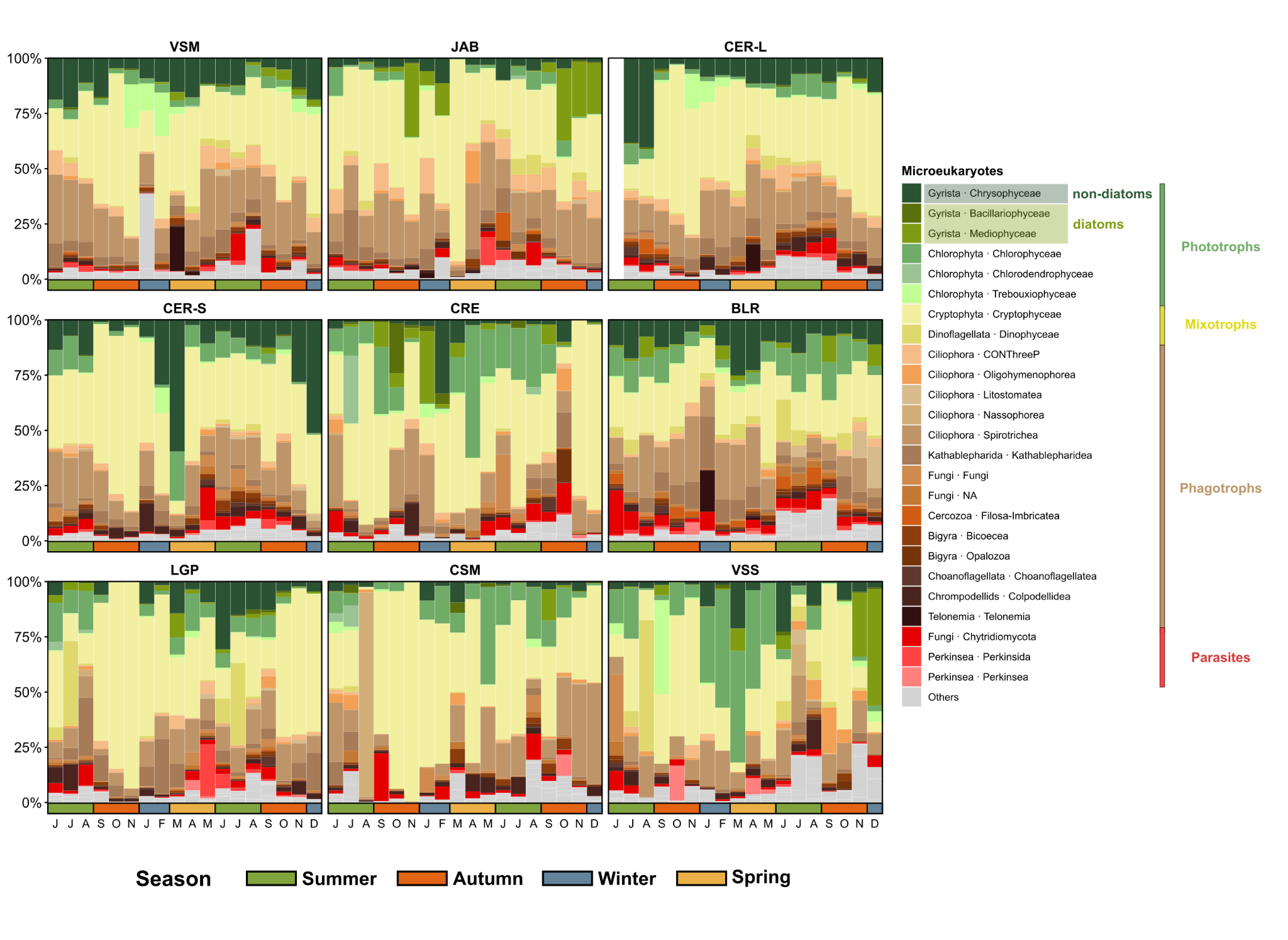

**Fig. S5: Temporal variation of the diversity indexes of microeukaryote communities**

**A-B:** Richness and Shannon diversity index over the 18 months (540 days) for the total ASVs. **C-F:** Richness of ASVs assigned to phototrophs (**C**), mixotrophs (**D**), phagotrophs (**E**), and parasites (**F**).

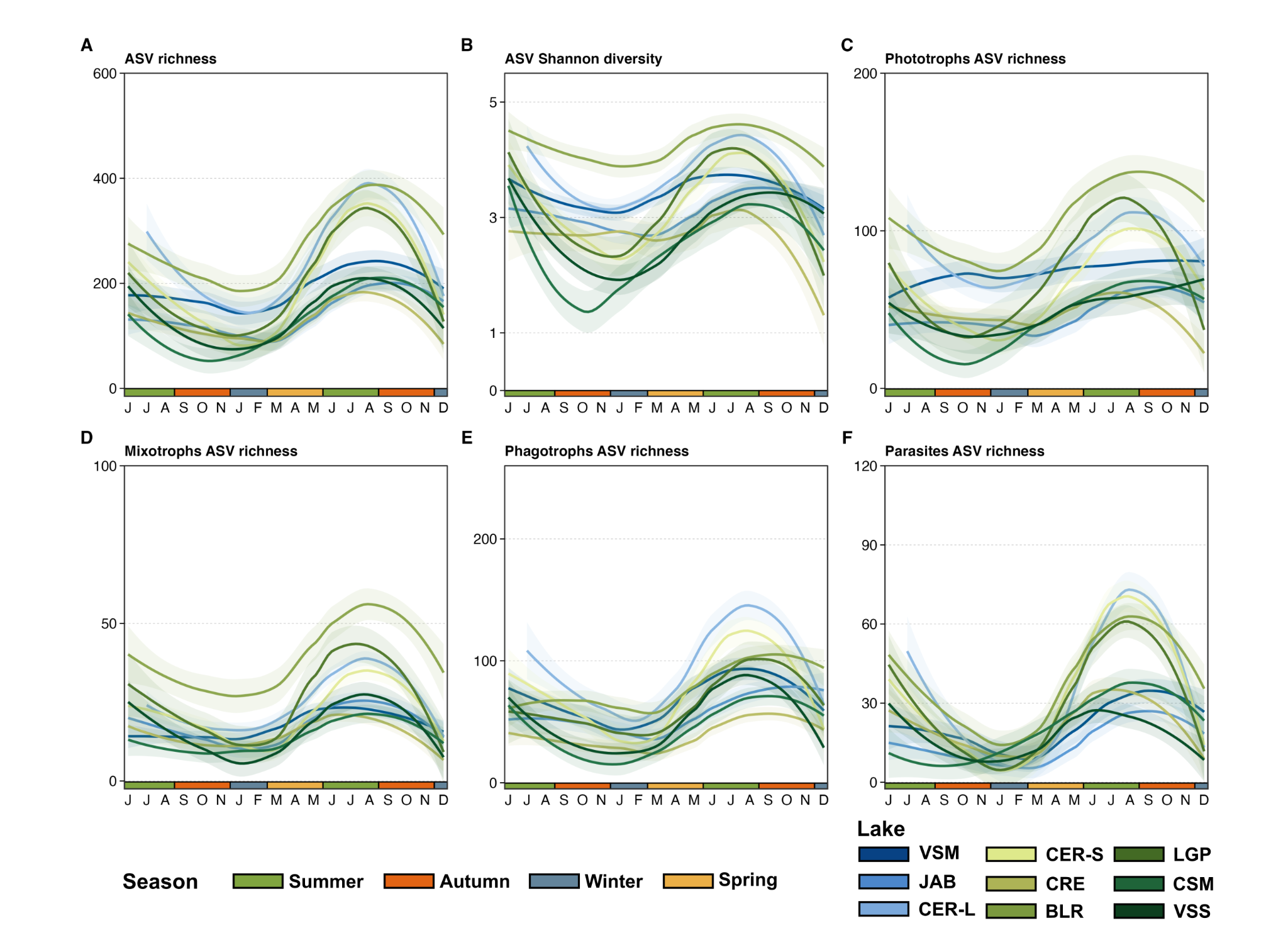

**Fig. S6: PCoA plots of the microeukaryote communities**

PCoA (based on Bray-Curtis dissimilarity, with all ASVs) from each lake are displayed in individual panels, with identical axis coordinates, and ordered according to their 18-month averaged Chl*a* concentration (from left to right, then from top to bottom). Seasons are colored and delimited by polygons representing the maximal area delimited by the sample’s coordinates.

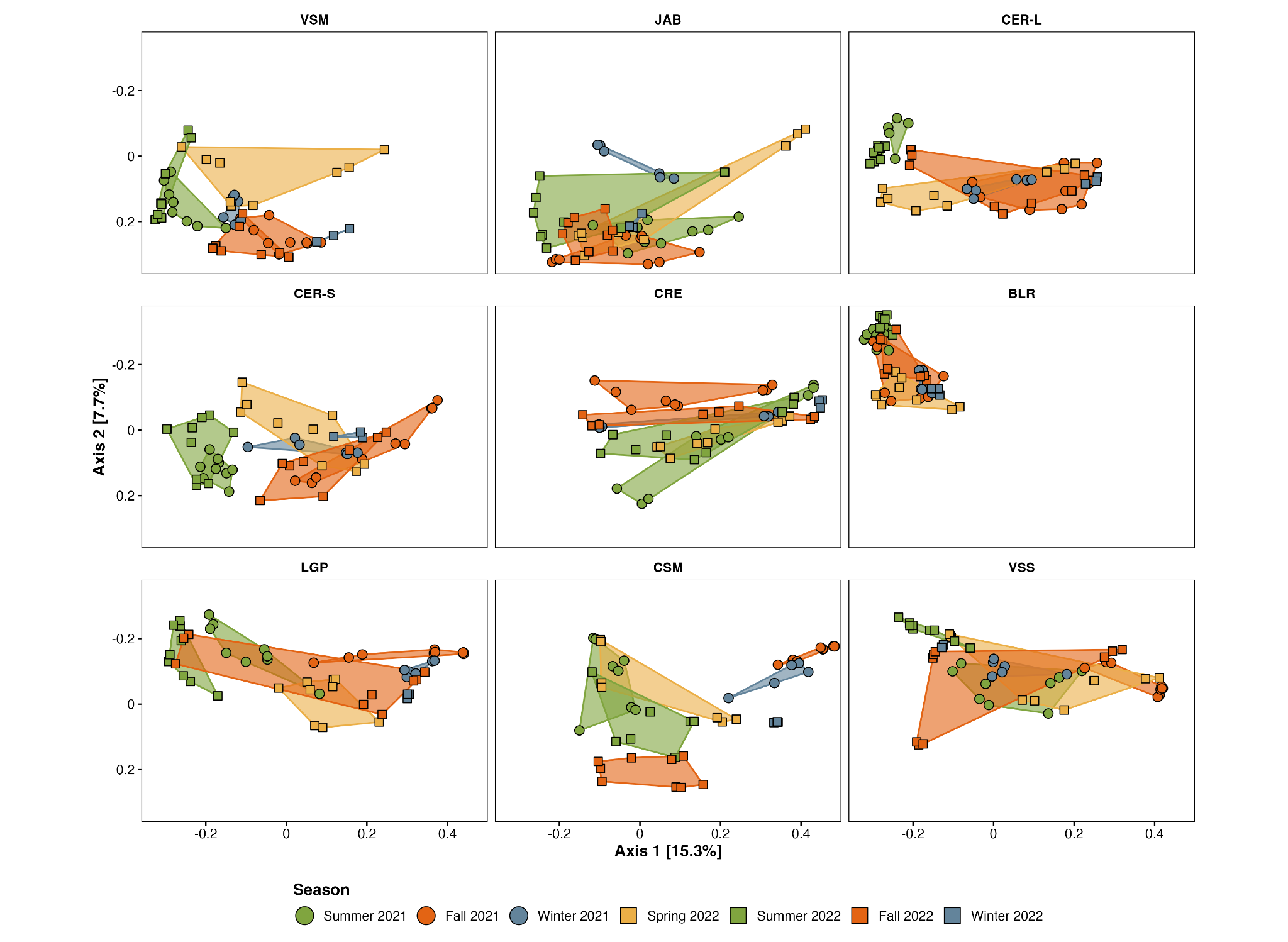

**Fig. S7: PCoA plots of the microeukaryote communities based on trophic modes**

PCoA (based on Bray-Curtis dissimilarity) based on ASVs classified as phototrophs (**A**), mixotrophs (**B**), phagotroph (**C**) and parasites (**D**) from each lake are displayed in individual panels. For each trophic mode the panels are displayed with identical axis coordinates and ordered according to their 18-month averaged Chl*a* concentration (from left to right, then from top to bottom). Seasons are colored and delimited by polygons representing the maximal area delimited by the sample’s coordinates.

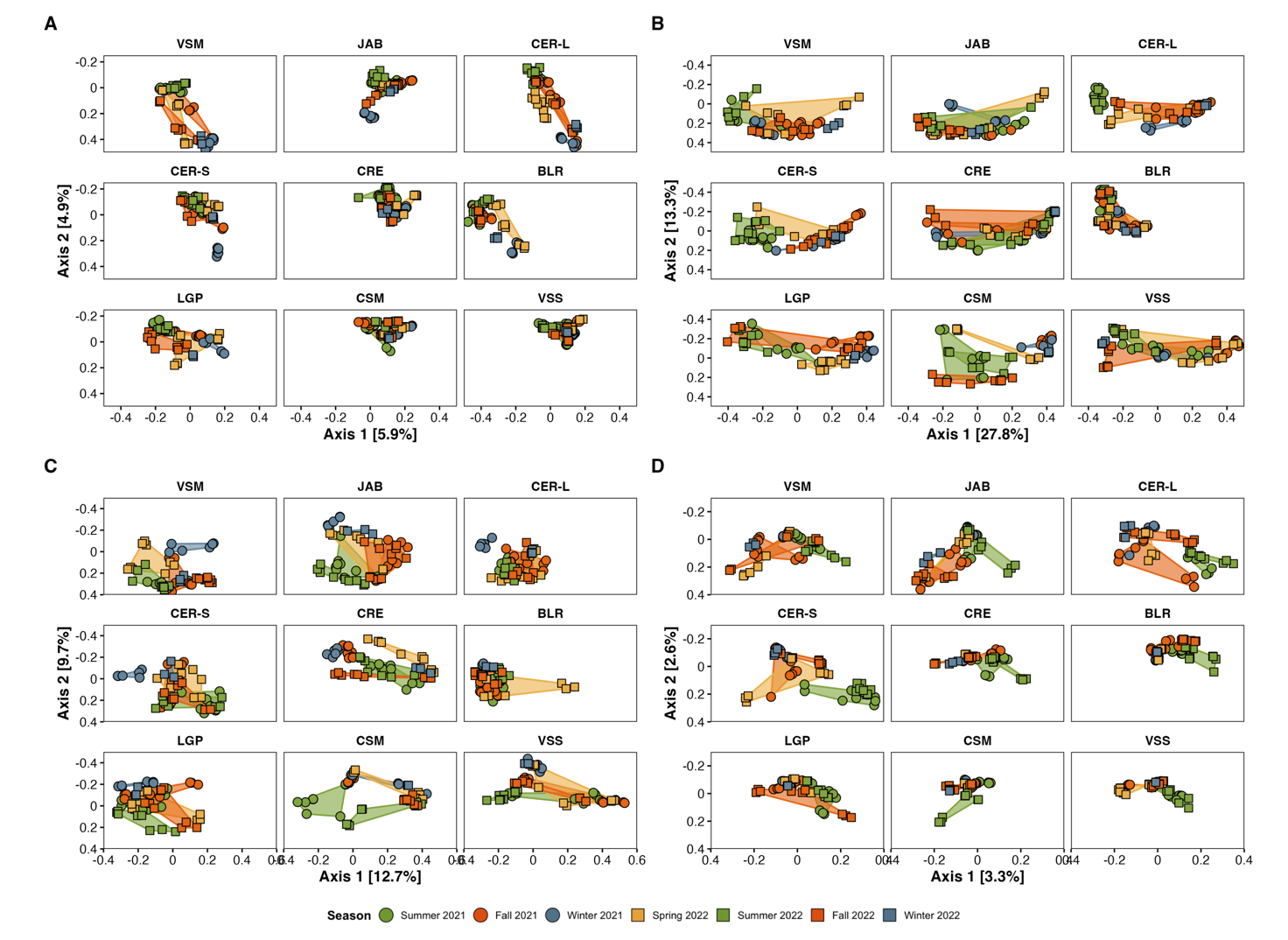
